# First Genome-Scale Metabolic Modeling of *Brucella abortus* Predicts Pathogen Gene Essentiality and New Drug Target

**DOI:** 10.1101/2023.10.19.563124

**Authors:** Bruno Ribeiro Pinto, Filipe Vilaça Guimarães de Oliveira, Miquéias Fernandes, Alex Jerônimo Ranieri Lima, Tiago Antônio de Oliveira Mendes

## Abstract

Bovine brucellosis, a globally widespread disease, imposes substantial economic burdens on livestock production. The pathogen, *Brucella abortus*, has a particular affinity for infecting cattle and can also impact humans, potentially posing a public health risk in regions where the disease persists. To gain a deeper understanding of the pathogen’s biology and its interactions with the host, Pathogen-specific Genome-Scale Metabolic Models offer valuable insights. They aid in identifying novel pharmacological targets and biomarkers, which can inform innovative brucellosis control strategies. In this study, we developed and validated the first Genome-Scale Metabolic Model of *B. abortus* 2308, named iBP932, encompassing 932 genes, 1,140 reactions, and 999 metabolites. Furthermore, iBP932 capability to predict potential drug targets was demonstrated.

## Introduction

*Brucella*, a genus of Gram-negative bacteria, is classified within the α-proteobacterial class (Batut et al, 2004). It is characterized as an aerobic coccobacillus, non-encapsulated, non-motile, facultatively intracellular, and urease-positive (Corbel and Hendry, 1985). The *Brucella* genus shelters around nine species, each pathogenic to distinct hosts (Moreno and Moriyón, 2002). Certain species within the genus exhibit positive oxidase, catalase, and superoxide dismutase activity (Araj, 2010). The presence of virulence factors like an unusual lipopolysaccharide, a type IV secretion system, cyclic β-1-2-glucans (CβG), Brucella virulence factor A (BvfA), and the BvrR/BvrS system are characteristics of the Brucella genus (Glowacka et al, 2018).

Among the members of the *Brucella spp*., *Brucella abortus* stands out as the etiological agent of bovine brucellosis. This is a worldwide spread disease with a high economic impact on livestock production, causing abortion, infertility, and weight loss in cattle (Gul and Khan, 2007). *Brucella spp.* possess the ability to adapt to novel hosts and can readily transmit to humans through direct contact with infected animals, as well as through the consumption of contaminated milk and its byproducts, thereby warranting attention in terms of public health concerns. Effective management of this disease entails animal vaccination, accurate diagnosis of infected animals, and culling of confirmed positive cases (Khurana et al, 2018).

The metabolic activities of intracellular pathogens such as *B. abortus* play a pivotal role in their virulence and ability to survive within the challenging intracellular environment. Gaining insights into these metabolic processes occurring within the host can facilitate the exploration of pathogen biology and identification of potential drug targets for effective control of the infection (Best and Kwaik, 2019). In this context, Genome-Scale Metabolic Models (GSMM) serve as a valuable knowledge base for understanding the intricate metabolic pathways of the pathogen. These models can be analyzed *in silico* using diverse Systems Biology approaches, enabling comprehensive investigations of the pathogen’s metabolism. One such approach is Flux Balance Analysis (FBA), which leverages the pathogen’s GSMM to make predictions about phenotypes, gene essentiality, and growth under various conditions (Teusink et al, 2010).

Pathogen-specific GSMMs can be used to study the pathogen metabolism, pathogen-host interactions, antibiotic resistance, virulence factors metabolism, biomarker research, and analysis of putative essential genes, which can be explored as new drug targets to design new therapeutic strategies. Furthermore, these data can be integrated with omics data to produce context-specific models (Sertbas, and Ulgen, 2020). While the central carbon metabolism of *B. abortus* has been extensively investigated (Barbier et al., 2017), the molecular mechanisms underlying infection remain poorly understood, and currently, there is a lack of a GSMM for *B. abortus.* In this study, we constructed the first GSMM of *B. abortus* by utilizing the existing genomic information, as well as metabolic data sourced from public databases and literature. Experimental data on gene essentiality, as well as information regarding carbon and nitrogen sources, were employed for model validation. Furthermore, the model was utilized to identify potential drug targets that exhibit broad-spectrum efficacy and do not share homology with human proteins.

## Methods

### Reconstruction of the metabolic network

To initiate the draft reconstruction of the GSMM of *B. abortu*s, a homology reconstruction process was carried out employing the RAVEN Toolbox version 2.0 (Wang et al., 2018), as illustrated in Figure 1. Initially, protein sequences extracted from the genomes of *B. abortus* 2308 (GenBank assembly accession: GCA_000054005.1) and the related organism *Sinorhizobium meliloti* 2011 (GenBank assembly accession: GCF_000346065.1) were utilized as input for the homology reconstruction process. For the draft homology reconstruction, the GSMM of *S. meliloti* was employed as a reference (diCenzo et al., 2020). The biomass equation of *S. meliloti* was modified to suit the *B. abortus* GSMM, considering their close phylogenetic relationship regarding differences in lipids and lipopolysaccharide (LPS) composition (Cardoso et al., 2006). Inclusion of cofactors and vitamins in the model was based on the comprehensive study by Xavier et al. (2017), which identifies essential cofactors and vitamins universal to prokaryotes. As the LPS of *B. abortus* differs from that of *S. meliloti*, it was incorporated into the biomass equation along with its corresponding biosynthetic pathway, as described by Cardoso et al. (2006). The Monomethyl phosphatidylethanolamine (MMPE) and sulfolipid sulfonoquinovosyl diacylglycerol (SL) were excluded from the biomass equation of *B. abortus*, as these lipids are not present in this organism (Sohlenkamp and Geiger, 2016). The proportions of amino acids and nucleotides were determined by extrapolating from the nucleotide and amino acid compositions of the predicted proteins encoded by the *B. abortus* genome (Zorrila and Kerkhoven, 2021). The final biomass equation of *B. abortus* model is composed of amino acids, nucleotides present in DNA and RNA, LPS, peptidoglycan, phospholipids, fatty acids, inorganic ions, cofactors and vitamins (Supplementary File S2).

**Figure 1.**
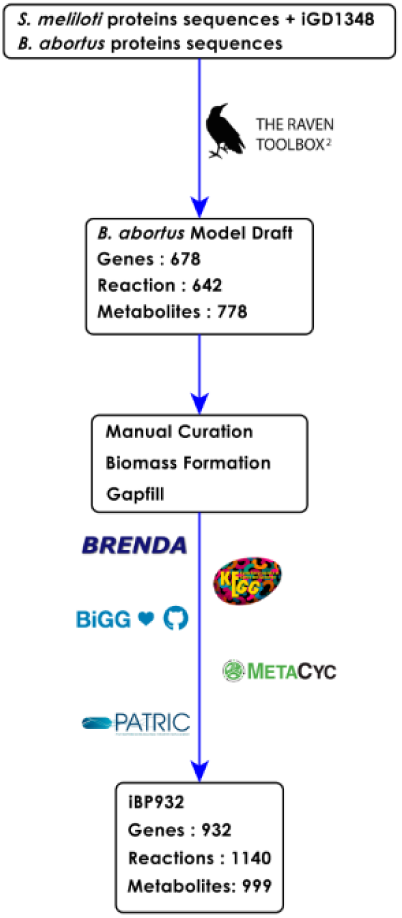
iBP932 reconstruction summary. In the initial phase, the *S. meliloti* model, iGD1348, and protein sequences, along with *B. abortus* proteins, were employed to draft a reconstruction through the utilization of the RAVEN Toolbox 2.0. Subsequently, following the draft generation, the formulation of a biomass equation and a manual curation process ensued, drawing upon information available in databases and the existing literature. Upon completion of the curation process, the final model, denoted as iBP932, was established.

Following the GSMM draft generation, an extensive manual curation process was undertaken to incorporate additional reactions, metabolites, and establish accurate Gene-Protein-Reactions (GPR) associations. This curation process involved leveraging the genomic annotation of *B. abortus* as well as utilizing several databases, including KEGG (Kanehisa et al., 2006), Metacyc (Caspi et al., 2020), BiGG (King et al., 2016), and PATRIC (Davis et al., 2020). To access the model functionality, the growth rate was predicted *in silico* and compared with experimental data. The growth rate from experimental data, was accessed based on the growth of *B. abortus* in a *Brucella* synthetic media. And the growth rate was calculated based on the growing curve. (Viadas et al., 2010). The *Brucella* synthetic media was reconstructed *in silico* and the uptake rate of each component was set to 1 mmol gDW^-1^ h^-1^. The description of the growth rate calculations and detailed media composition are described in Supplementary Material File S1 and S2. All constraint-based flux analyses and tests conducted during the draft generation phase were implemented using Constraint-Based Metabolic Modeling in Python (COBRApy) version 0.25.0 (Orth et al., 2010; Mahadevan and Schilling, 2003). The finalized and curated model was designated as iBP932.

### Model validation Gene Essentiality

To validate the reconstruction of iBP932, an *in silico* analysis of gene essentiality was conducted. The predictions obtained were compared to previous in vitro investigations that identified genes crucial for survival within macrophage cells (Bonomi et al., 2010; Gao et al., 2013; Sternon et al., 2018; Tian et al., 2019; Révora et al., 2020). Additionally, the gene essentiality dataset available in the BioCyc database for *B. abortus* 2308 (BioCyc ID: MIX2B4Q-7) was utilized. This dataset is based on the research published by Sternon et al. (2018) and encompasses growth conditions using the 2YT complex medium as well as macrophage infection experiments.

To achieve accurate gene essentiality results, the 2YT complex medium growth was modeled *in silico* following the methodology described by Marinos et al, 2020. This approach considers the comprehensive composition of amino acids, nucleotides, vitamins, and ions present in the complex medium components such as yeast extract and tryptone. The model incorporates exchange reactions to account for the metabolites derived from the medium, and rather than employing uptake rates, the concentration of individual components is utilized (see Supplementary Material File S2).

Under *in vivo* conditions, during the initial stages of infection, *Brucella spp*. establishes residence within the phagosomal compartment of host macrophages and endures hostile environmental factors such as reactive oxygen intermediates, acidic pH, and nutrient scarcity. In response to nutrient deprivation, bacteria enter a physiological state known as the stationary phase, which allows for the maintenance of metabolism and cell viability (Roop II et al., 2003). During macrophage infection, *B. abortus* induces a metabolic shift known as the Warburg effect in the host cell, leading to increased glycolysis and the production of lactate, which is used as a carbon and energy source (Czyz et al., 2017). Thus, in the simulation of macrophage conditions (MFG), lactate was employed as the carbon source.

A study investigating the nitrogen sources available to *Mycobacterium tuberculosis* within human macrophages revealed the presence of amino acids, including glutamine, alanine, aspartate, glutamate, valine, leucine, and glycine. Among these, glutamine was identified as the primary nitrogen donor (Borah et al., 2019). Considering this study, glutamine and the other six amino acids, alanine, aspartate, glutamate, valine, leucine, and glycine, were constrained as nitrogen sources for the MFG modeling. To assess the impact of gene deletions, simulations were performed utilizing the COBRApy single gene knockout function. This enabled the investigation of the consequences resulting from the removal of specific genes within the metabolic network.

### Nitrogen and Carbon Sources Utilization

Prior investigations have demonstrated the ability of certain strains of *B. abortus* to utilize diverse carbon sources via the Biotype 100 carbon assimilation system (López-Merino et al., 2000). Additionally, biotyping systems have been employed to assess metabolic activity to differentiate various members of the *Brucella* species (Wong et al., 1992; Al Dahouk et al., 2010). To validate the reconstructed iBP932 model, an analysis was conducted to examine the utilization of metabolites as carbon and nitrogen sources. This analysis was based on the methodology developed by Al Dahouk et al. (2010). Metabolites associated with high and moderate metabolic activity in *B. abortus* were selected for validation purposes. A chemically-defined minimal medium was utilized, with the exchange of lactate and glycerol for D-glucose as the carbon source and ammonia as the nitrogen source (Gerhardt and Wilson, 1948). In this analysis, D-glucose was substituted with other carbon sources to be tested, while ammonia was replaced with alternative nitrogen sources. FBA was subsequently employed to evaluate the *in silico* growth under these conditions.

### Gene-Centric Identification of Drug Targets for iBP932

The identification of potential drug targets was based on the methodology established by Cesur et al. (2020). This approach involved a two-step process. Initially, the essentiality of each gene in the metabolic network was determined by constraining the model to specific growth conditions. In the second step, genes exhibiting no homology to humans were selected and tested for the following criteria, druggability, potential virulence factors and broad-spectrum (Cesur et al., 2020). Gene essentiality was assessed using the single-gene deletion function from COBRApy under two distinct conditions: simulating Human Blood Fluids (HBF) (Hadi and Marashi, 2014) and the macrophage intracellular environment using the same gene essentiality constraints applied in model validation. The HBF condition was defined by 77 metabolites, which were used as constraints corresponding to the uptake reactions present in the iBP932 model. During HBF simulations, the uptake rate of all media components was set to 10 mmol gDW-¹ h-¹ with the biomass equation serving as the objective function in order to predict the growth rate during FBA.

### Druggable Targets Analysis

In order to mitigate the selection of drug targets that could pose harm to the human host, we employed the following strategy. Initially, protein sequences corresponding to essential genes predicted by iBP932 were subjected to BLASTp analysis against human protein sequences (GenBank assembly accession: GCA_000001405.29) using an expected value (E-value) cut-off of 1x 10-4. Proteins encoded by genes with E-values exceeding the cut-off and present in the essential gene list were considered devoid of homology to humans and selected for further analysis. Next, we assessed the druggability of the identified targets that were non-homologous to humans but predicted as essential to *B. abortus*. Druggability refers to the capacity of a target to bind tightly to drug-like compounds. To evaluate druggability, we utilized the Drugbank database (Wishart et al., 2008). The non-homologous proteins were subjected to a BLASTp search against the known targets in the Drugbank database, adopting an E-value cut-off of 1x10-25.

### Potential Virulence Factor Analysis

Virulence factors encompass a diverse array of molecules such as toxins, enzymes, lipopolysaccharides, as well as cell surface structures like capsules, LPS, glycoproteins, and lipoproteins. These molecules are produced by microbial pathogens to subvert the host’s defense mechanisms and establish an infection (Leitão, 2020). To identify drug targets that may also serve as anti-virulence agents for more effective drug therapy, we investigated whether any of the druggable proteins identified could be potential virulence factors. For this purpose, we employed the virulence-factor database VDFB (Chen et al., 2016) and conducted a BLASTp analysis against the database, considering as valid hits which presented an E-value cut-off of 1x10-4, a bit score of 100, and an identity threshold of 65% (Cesur et al., 2020).

### Broad-Spectrum Analysis

To assess the potential for broad-spectrum activity in drug target candidates, which can be employed for the treatment of co-infections or diverse infections, we performed a broad-spectrum analysis. This analysis involved a homology search against various infectious bacteria utilizing the comprehensive dataset of PBIT (Pipeline Builder for Identification of Drug Targets) web browser (Shende et al., 2017). Specifically, we executed a BLASTp search against protein sequences of pathogenic organisms, applying an E-value cut-off of 1x10-5, a bit score of 100, and a sequence identity threshold of 35%. A protein was considered a broad-spectrum target if it was identified in at least 40 diverse pathogenic strains (Cesur et al., 2020) Figure 2.

**Figure 2.**
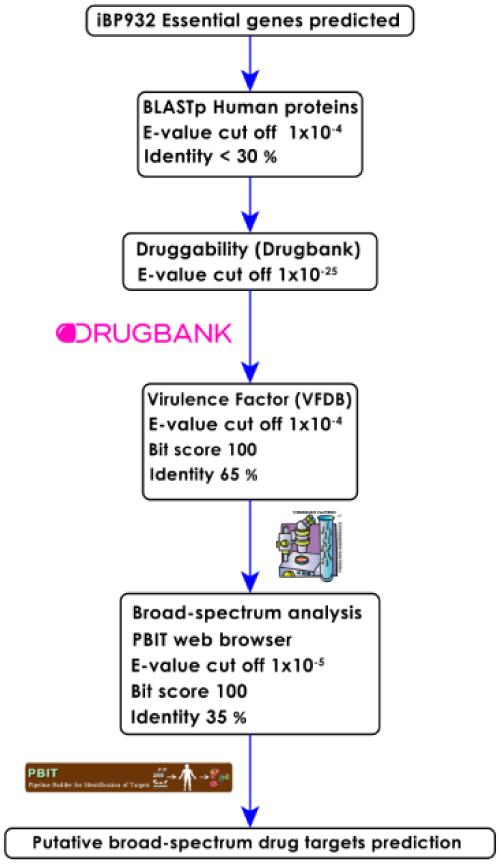
Drug Targets selection pipeline.

## Results and Discussion

### Model Reconstruction

The resulting model reconstruction of the *B. abortus* 2308 was named iBP932, which consists of 1,140 reactions, 999 metabolites, 932 genes, and three cellular compartments: cytoplasm, extracellular, and the periplasm. The total of 932 genes included in the iBP932 represents 26.7 % of the genes present in the bacterial genome, which codes for enzymes and transporters of a total of 3.034 genes distributed in a genome size of 3.2 Mb. Among the reactions included in this model, 28 are non-gene associated. These reactions either occur spontaneously or have been incorporated into the model to fulfill the requirement for the production of biomass components. A comprehensive overview of the model is displayed in Table 1.

**Table 1.** Features of *B. abortus* genome-scale metabolic model iBP932.

| Features of iBP932 | Number |
| --- | --- |
| Total genome size (bp) | 3.278.307 |
| Open reading frames (ORFs) | 3.034 |
| Gene Included in the model (GPR) | 932 |
| Model Reactions | 1.140 |
| Model Reactions Without GPR | 36 |
| Purely Metabolic Reactions | 862 |
| Transport Reactions | 139 |
| Metabolites | 999 |

The establishment of a precise biomass equation is a pivotal requirement in the process of reconstructing metabolic models. This equation bears a direct correlation with the accuracy of predicting gene essentiality, which holds great significance in the context of pathogen GSMMs. In this study, we opted for the adaptation of the biomass equation from the GSMM of *S. meliloti*, primarily due to the close phylogenetic relationship between this organism with *B. abortus*. Furthermore, in order to generate a biomass equation that is more specific to *Brucella*, adjustments were made to account for the lipid composition and other virulence factors such as LPS. This refined biomass equation, tailored to *Brucella*, holds the potential to serve as a template for the development of GSMMs for other *Brucella* species. Moreover, it also lays the foundation for further refinement and enhancement of the equation itself. The iBP932 *in silico* growth rate in the *Brucella* synthetic media was 0.058 h^-1^ predicted using FBA and the biomass equation as objective function. Compared with experimental data, 0.052 h^-1^, the model shows a very close value in the growth rate. These results validate the iBP932 *in silico* growth prediction. The construction of a functional GSMM for *B. abortus* was made possible through the integration of information derived from genomic annotation, scientific literature, and biological databases. Further investigations into the biochemistry and physiology of *B. abortus* have the potential to provide valuable insights that can be utilized to enhance and refine the model, advancing our understanding of this pathogen’s metabolic capabilities.

### Gene Essentiality Analysis

In the context of GSMMs for pathogens, the analysis of gene essentiality plays a critical role in model validation, as the predictions inferred from the model can contribute to the identification of feasible antimicrobial drug targets. In the dataset obtained from BioCyc (BioCyc ID: MIX2B4Q-7), based on a comprehensive Tn-seq study (Sternon et al., 2018) revealed that 476 single gene knockouts of *B. abortus* resulted in no growth in 2YT culture media. Among these studied genes, a total of 275 are present in the iBP932 model. To assess the accuracy of iBP932 in predicting gene essentiality, single *in silico* gene knockout simulations were performed in 2YT complex media, and the results are presented in Figure 3. A comparison between the gene essentiality predictions of iBP932 and experimental data showed a remarkable accuracy of 79%. This value falls within an acceptable range and surpasses the accuracy of other published pathogen GSMMs, such as *Acinetobacter baumannii* with 72% accuracy (Kim et al., 2009), *Streptococcus oralis* with 71-76% accuracy (Jensen et al., 2020), and *Mycobacterium tuberculosis* with 71% accuracy (Kavvas et al., 2018).

**Figure 3.**
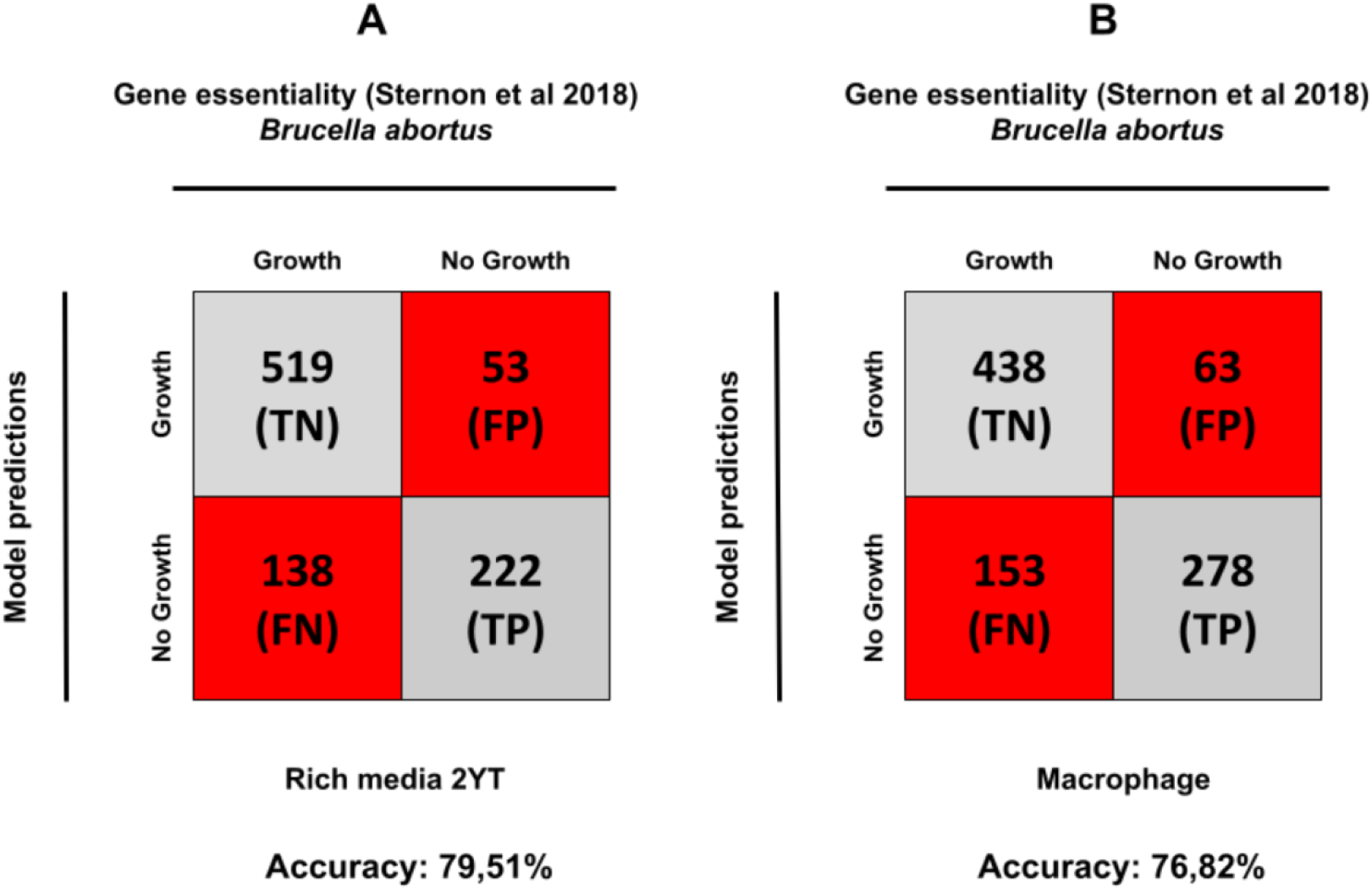
Gene essentiality comparison between iBP932 and experimental data. **(A)** When iBP932 is compared with the experimental data of Sternon et al 2018, 2YT rich media condition there is a 79% accuracy with gene essentiality. **(B)** When comparing with macrophage *in vivo* condition the gene essentiality accuracy of iBP932 is 76% also based on the Sternon et al 2018 experimental data.

Among the instances of false positive results in gene essentiality analysis (genes that are experimentally essential and computationally not essential), a notable observation pertains to genes associated with the Tricarboxylic Acid (TCA) cycle. When these genes are computationally deleted in silico, the TCA intermediates can still be produced through alternative reactions present in the model, particularly catabolic reactions involved in the degradation of amino acids. It is important to note that the computational model predicts a steady-state condition, wherein the majority of reactions governing growth prediction are biosynthesis reactions. In contrast, catabolic reactions primarily occur when the cell requires a breakdown of molecules for energy generation or nutrient utilization.

In the steady-state condition of a GSMM, the deactivation of any reaction triggers the network to reconfigure itself by adopting alternative pathways to achieve the optimal solution for the given objective function. This observation highlights the potential benefit of integrating transcriptomic data into context-specific modeling approaches, as it can provide valuable constraints to better align the model with experimental conditions. In the study conducted by Sternon et al. (2018), the gene essentiality of *B. abortus* was assessed specifically in macrophages. In order to evaluate the accuracy of gene essentiality predictions of the iBP932 model, we incorporated the MFG growth conditions into the model. The comparison between the model predictions and the experimental data yielded a commendable accuracy of 76%. This demonstrates the reliability and utility of the iBP932 model in predicting gene essentiality under the specific growth conditions encountered in macrophages.

The study by Sternon et al. underscores the requirement for the presence of complete biosynthetic pathways for histidine and pyrimidine in order to support growth. To examine the essentiality of these pathways under macrophage condition, we conducted simulations within the iBP932 model by systematically applying gene knockouts and deactivating individual reactions within the biosynthetic pathways for histidine and pyrimidines. The results revealed that the iBP932 model also relies on the presence of these intact pathways to sustain *in silico* growth Figure 4. This finding further supports the critical role of these biosynthetic pathways in supporting the metabolic requirements of *B. abortus*.

**Figure 4.**
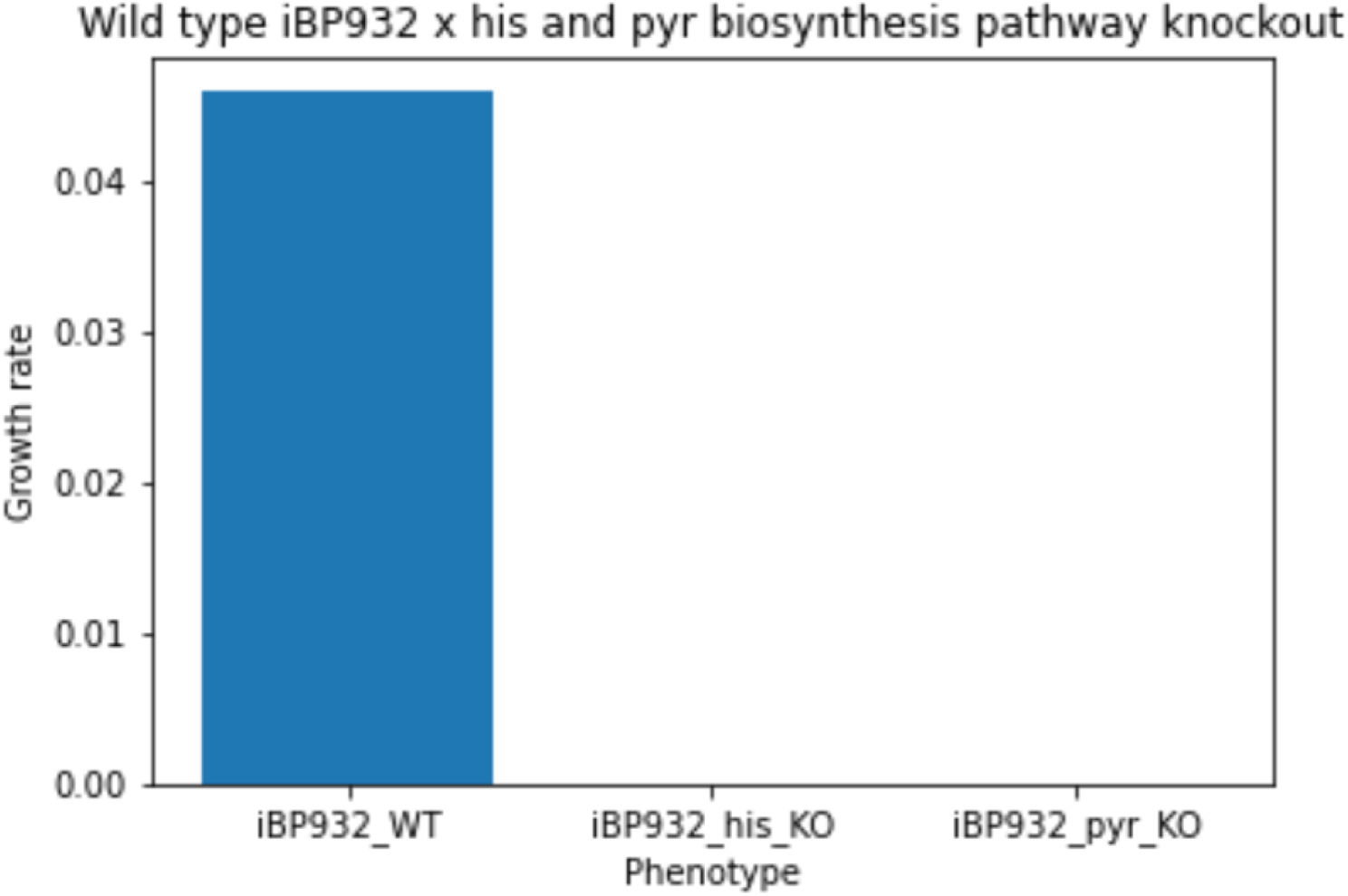
The histidine biosynthesis pathway in iBP932 comprises ten reactions: ATPPPRT (hisZ and hisG), PRATPP (hisE), PRAMPC (hisI), PRMICI (hisA), IG3PS (hisH and hisF), IGPDH (hisB), HSTPT (hisC), HISTP (hisN), HISTD, and HISND (hisD). The pyrimidine biosynthesis pathway, comprising the reactions ASPCT (pyrB), DHORTS (pyrC), DHORDi (pyrD), ORPT (pyrE), and OMPDC (pyrF). Detailed information regarding these reactions can be accessed in the iBP932 Excel format provided in the supplementary material.Following the in silico knockout of all histidine of pyrimidine biosynthesis genes, it is predicted that no growth would occur under MFG conditions, consistent with the findings of Sternon et al. in their 2018 in vitro study. This observation underscores the indispensable requirement for a fully functional histidine and pyrimidines biosynthesis pathway to ensure survival within macrophages.

### Utilization of Carbon and Nitrogen Sources by iBP932: in silico and in vitro conditions

To assess the model’s validity in simulating growth across diverse carbon and nitrogen sources, we employed the metabolic test devised by Al Dahouk et al. in 2010. This test leverages enzyme activities and the ability to metabolize various substrates as carbon and nitrogen sources to phenotypically differentiate *Brucella spp*. In our study, we evaluated a total of 27 metabolites that were experimentally confirmed to be utilized by *B. abortus*. Among these metabolites, 19 (of a total of 27) exhibited positive outcomes for biomass production when simulated using the iBP932 model, while 8 showed negative results. The metabolites yielding negative outcomes were D-glucose-L-cysteine, D-Talose, D-threitol, Gallic acid, L-cystine, L-carnosin, D-arabinose, and Guanidinosuccinic acid.

The identification of metabolites yielding negative results served as a starting point for refining the model. For instance, the metabolite D-threitol, which demonstrated unfavorable outcomes, was investigated based on the research conducted by Huang et al. in 2015. Huang et al. elucidated the catabolic pathway of this polyol in *Mycobacterium smegmatis*, involving four enzymes: D-threitol dehydrogenase (DthD), D-erythrulose kinase (Derk), and D-erythrulose-4-phosphate isomerase (DeIr1 or Der2) (Huang et al., 2015). To address this gap in the iBP932 model, a homology search was performed using BLASTp and InterproScan (Blum et al., 2020) to identify homologous enzymes within the *B. abortus* genome. The search focused on examining protein domains. Consequently, a gene encoding D-threitol dehydrogenase (BAB_RS23775) was identified in the *B. abortus* genome. For the D-erythrulose kinase, the transporter (BAB_RS23780) and two copies of genes encoding subunits of this enzyme (BAB_RS23755, BAB_RS23750, BAB_RS23770, and BAB_RS23765) were found. As for the D-erythrulose-4-phosphate isomerase, a single gene (BAB_RS28110) encoding this enzyme was identified in the *B. abortus* genome.

The metabolism of gallic acid has been extensively characterized in *Pseudomonas putida*, where enzymes encoded by the gal cluster play a pivotal role (Nogales et al., 2011). However, the metabolic pathway for gallic acid in *B. abortus* remains poorly understood. In the experiments conducted by Al Dahouk et al., all *Brucella* species exhibited significant metabolic activity towards gallic acid. To elucidate the metabolic pathway in *B. abortus*, a homology-based gene search was performed within the *B. abortus* genome targeting the gal cluster. As a result, two genes encoding the enzyme protocatechuate 3,4 dioxygenase were identified, which have been described in MetaCyc as part of the gallic acid biodegradation pathway II (BioCyc ID: GALLATE-DEGRADATION-I-PWY). These enzymes catalyze the initial step of gallic acid metabolism, as previously described by Sparnis and Dagley (Sparnis and Dagley, 1975). Additionally, two putative genes were found: one for galT, encoding a transporter for gallic acid, and another for galB, encoding the enzyme 4-oxalmesaconate hydratase. These findings enabled the careful incorporation of the transport and metabolic reactions of gallic acid into the model.

In the case of L-carnosine, a potential beta-ala his peptidase (BAB_RS16495) responsible for the conversion of L-carnosine into *β*-alanine and L-histidine was identified in the *B. abortus* genome. This enzyme was successfully included in the model to account for L-carnosine metabolism. Regarding the other metabolites that exhibited negative results for *in silico* growth, no information regarding their metabolism was discovered within the *B. abortus* genome. Despite extensive searches, no relevant genes or pathways associated with these metabolites were found.

After this meticulous curation process, it was observed that 22 out of the 27 (81%) simulated metabolites (Figure 5) allowed for the prediction of biomass production. The utilization of various carbon and nitrogen sources appears to be vital for the survival of microorganisms during the infection process, thereby contributing to a better understanding of the pathological mechanisms involved. The iBP932 model exhibits the ability to predict growth under different conditions, considering diverse sources of carbon and nitrogen utilization. The identification of negative results serves as a crucial reference point for further refinement of the model.

**Figure 5.**
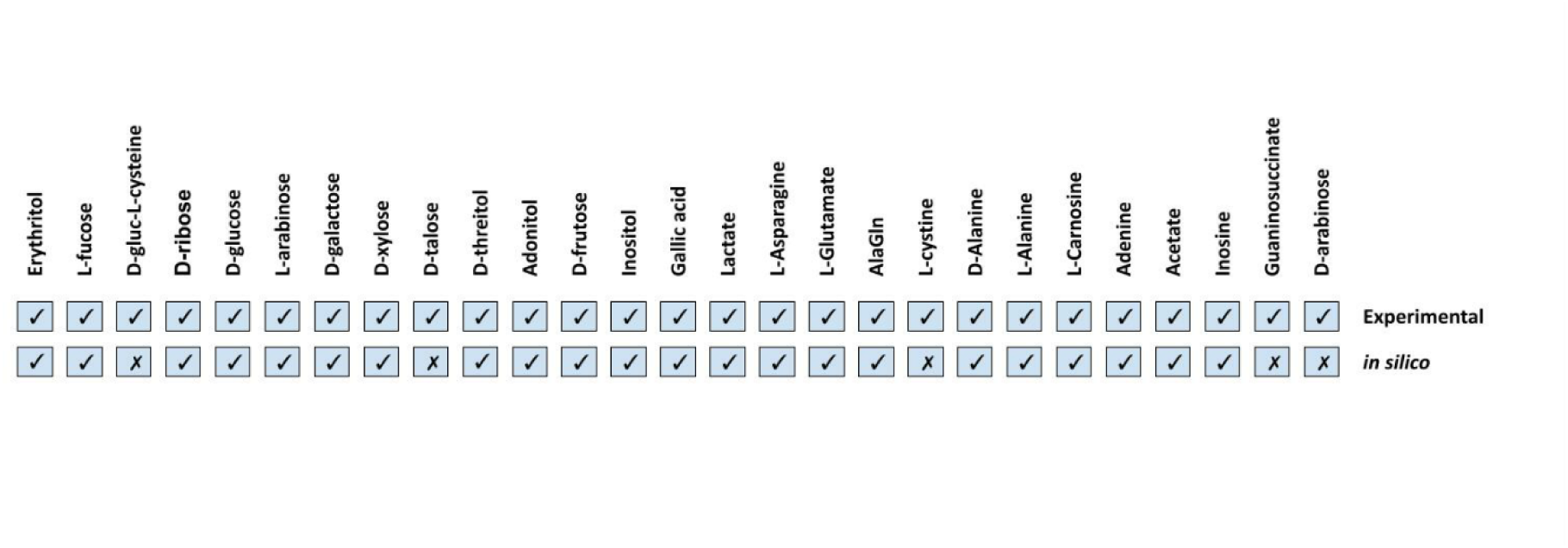
Carbon and nitrogen sources utilization validation. The results presents de 22 of 27 positive results of nitrogen and carbon utilization based on the Al Dahouk et al 2010 metabolic test, the ‘X’ in the boxes represent negative results, indicating that iBP932 cannot use that source to predict *in silico* growth and the ‘✓’ indicate the positive results comparing the *in silico* utilization of iBP932 of these substrates to predict growth a total of 81% accuracy.

### Essential gene analysis and drug targets selections of human blood fluid and macrophage conditions

The investigation of potential drug targets for two distinct conditions, human blood fluid (HBF) and MFG, resulted in the identification of 183 and 245 non-human homologous essential genes, respectively, within the iBP932 model (Supplementary material file 2). Notably, two distinct metabolic profiles emerged from this analysis. The MFG condition displayed a greater reliance on biosynthetic pathways involved in arginine, cysteine, histidine, aromatic, and branched amino acids, compared to the HBF condition. In particular, the HBF condition exhibited four essential genes associated with reactions in the biosynthetic pathway of aromatic amino acids, which are also vital in the MFG condition. These genes play a pivotal role in chorismate synthesis, highlighting the significance of chorismate within the iBP932 network. Chorismate is a critical metabolite required for the biosynthesis of aromatic amino acids, siderophores, and ubiquinone, underscoring its indispensability in various cellular processes.

Several crucial metabolic pathways were found to be common to both HBF and MFG conditions Figure 6. Notably, one such shared pathway is the biosynthesis of lipopolysaccharides (LPS), which holds significance in *Brucella* infection. LPS plays a pivotal role in the interaction between *Brucella* and the host, enabling the evasion of immune defense mechanisms and facilitating intracellular internalization (Cardoso et al., 2006). Given the criticality of LPS in *Brucella* infection, enzymes involved in LPS biosynthesis represent promising candidates for drug target exploration. Targeting these enzymes could impede *Brucella’s* ability to evade host defenses, thereby thwarting the establishment of infection.

**Figure 6.**
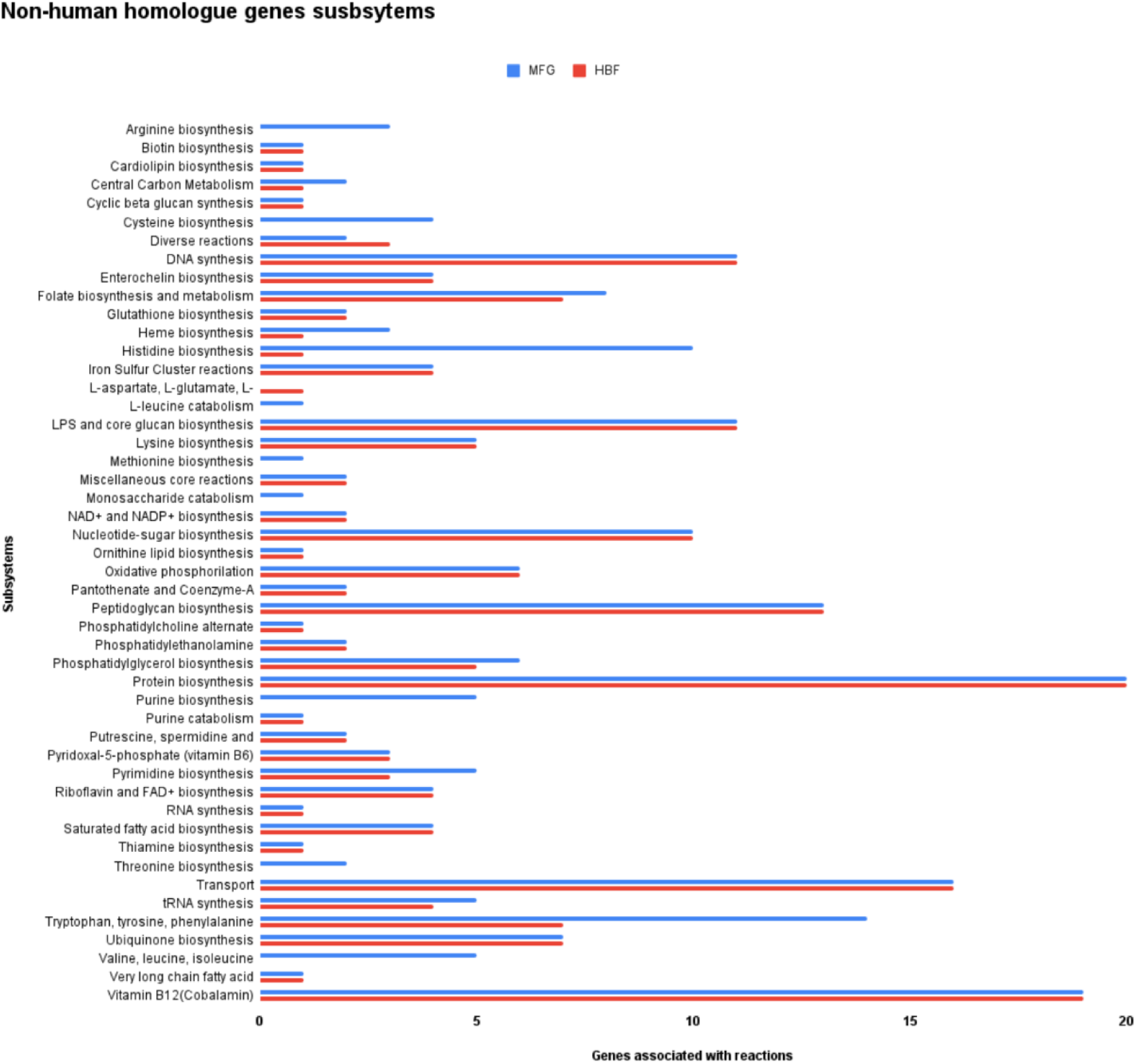
Comparison of the non-human homologue essential gene subsystem distribution between conditions HBF and MFG.

Vitamins play a critical role in maintaining metabolic processes, and adenosylcobalamin (B12) is particularly noteworthy. This vitamin binds to a diverse range of ligands, including proteins and nucleic acids, and is crucial for catalytic activities in B12-dependent enzymes (Gruber et al., 2011). The simulation involving the deletion of both RibH1 and RibH2 genes in the iBP932 model identified them as essential components. A previous investigation demonstrated that the double knockout of RibH1 and RibH2 genes, which encode two isoenzymes of lumazine synthase involved in riboflavin biosynthesis, resulted in *B. abortus* mutants incapable of surviving within cells and mice. These findings unveil the potential of targeting RibH1 and RibH2 genes for the treatment of brucellosis (Bonomi et al., 2010; Serer et al., 2014). Moreover, the enzyme riboflavin synthase, which catalyzes the final step of riboflavin biosynthesis, emerges as an additional promising drug target candidate. These studies underscore the vital role of vitamins, such as riboflavin, in the survival of Brucella and highlight their potential significance as targets for therapeutic interventions.

### Potential Drug Targets Identification

The non-homologous essential proteins were subjected to further analysis based on druggability, virulence factors, and broad-spectrum characteristics. Initially, druggability was assessed to identify proteins with similarities to known drug targets. For the two analyzed conditions, a total of 69 potential drug targets were identified for MFG and 47 for HBF (Table 2), with MFG encompassing all the targets identified in HBF. These targets shared common metabolic pathways involved in amino acid and vitamin metabolism, as well as peptidoglycan biosynthesis. Notably, 17 targets exhibited affinity with at least five drug-like molecules. Among the identified targets, four significant matches were found to interact with approved antimicrobials. Dihydropteroate synthase demonstrated an affinity for ten approved sulfonamides, while cycloserine, tigecycline, and clindamycin interacted with three respective putative drug targets: D-alanine-D-alanine ligase, 30S ribosomal protein S13, and 50S ribosomal protein L1 (Supplementary Material File S3). Subsequently, the selection of druggable targets was followed by the prediction of virulence targets using the complete dataset from VFDB. Only six targets met the criteria, including the genes dhdB (Isochorismatase), *lpxA* (UDP-N-acetylglucosamine acyltransferase), *lpxC* (UDP-3-O-acyl-N-acetylglucosamine deacetylase), *kdsB* (3-deoxy-manno-octulosonate cytidylyltransferase), *kdsA* (2-dehydro-3-deoxyphosphooctonate aldolase), and *FabZ* (3-hydroxyacyl-ACP dehydratase). These identified factors participate in various pathways, such as aromatic amino acid biosynthesis, LPS and core glucan biosynthesis, nucleotide-sugar biosynthesis, and saturated fatty acid biosynthesis. Of particular interest, the *dhdB* gene codes for the enzyme isochorismatase in the iBP932 model, which catalyzes the conversion of isochorismate to 2,3-dihydro-2,3-dihydroxybenzoate and pyruvate. Proteome analysis has indicated isochorismatase as a putative drug target candidate for *Brucella melitensis* (Rahman et al., 2021).

**Table 2.**
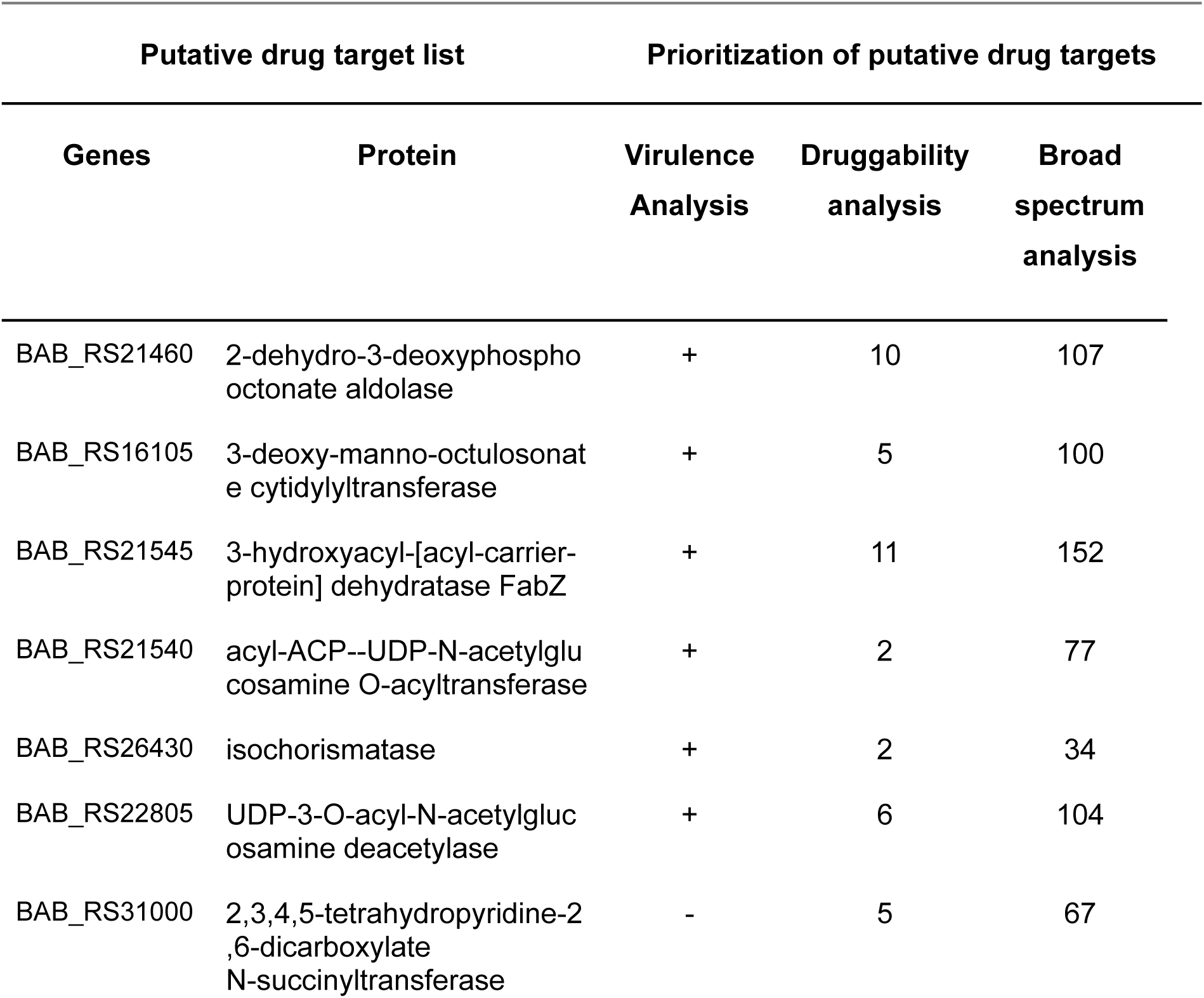
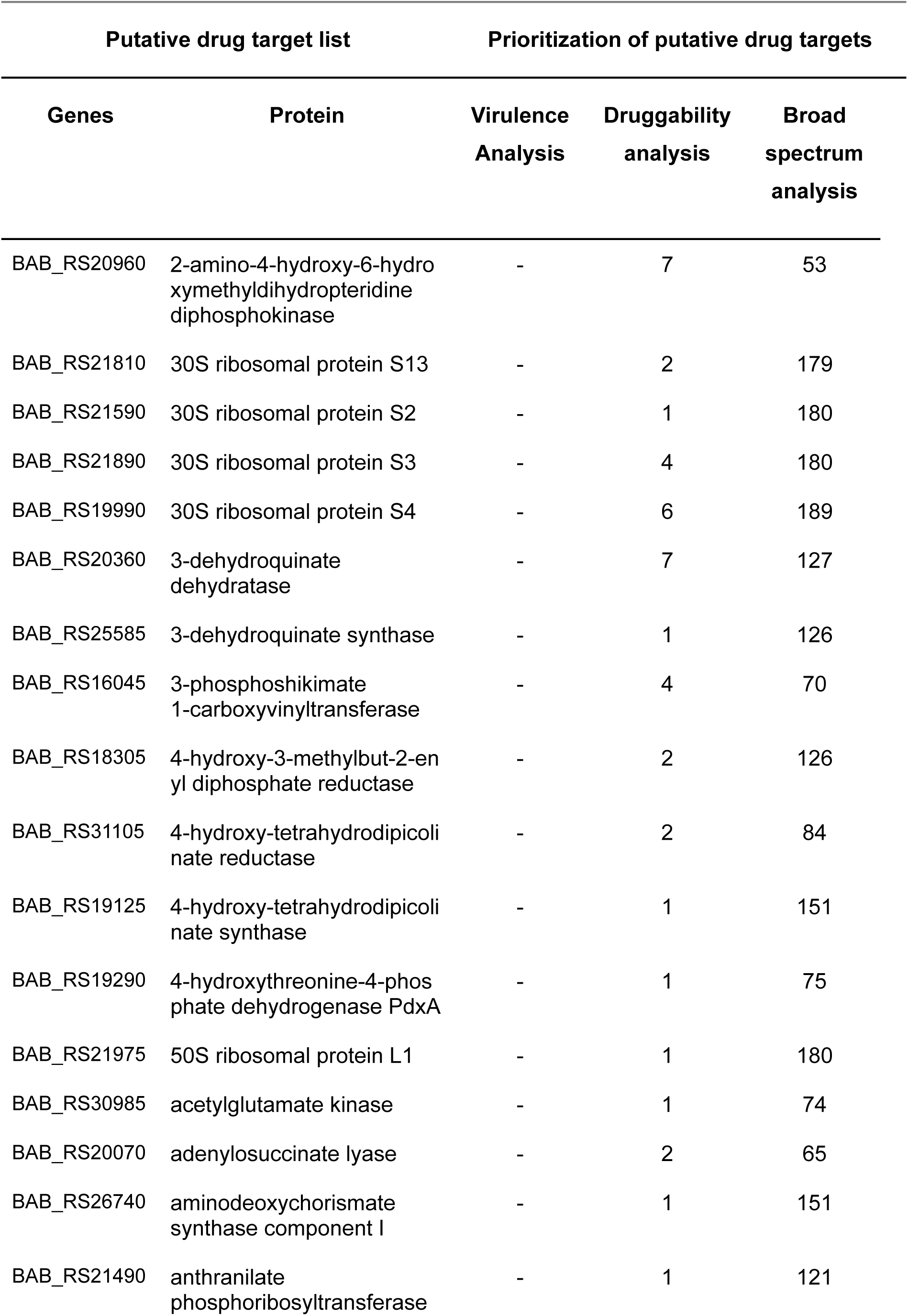
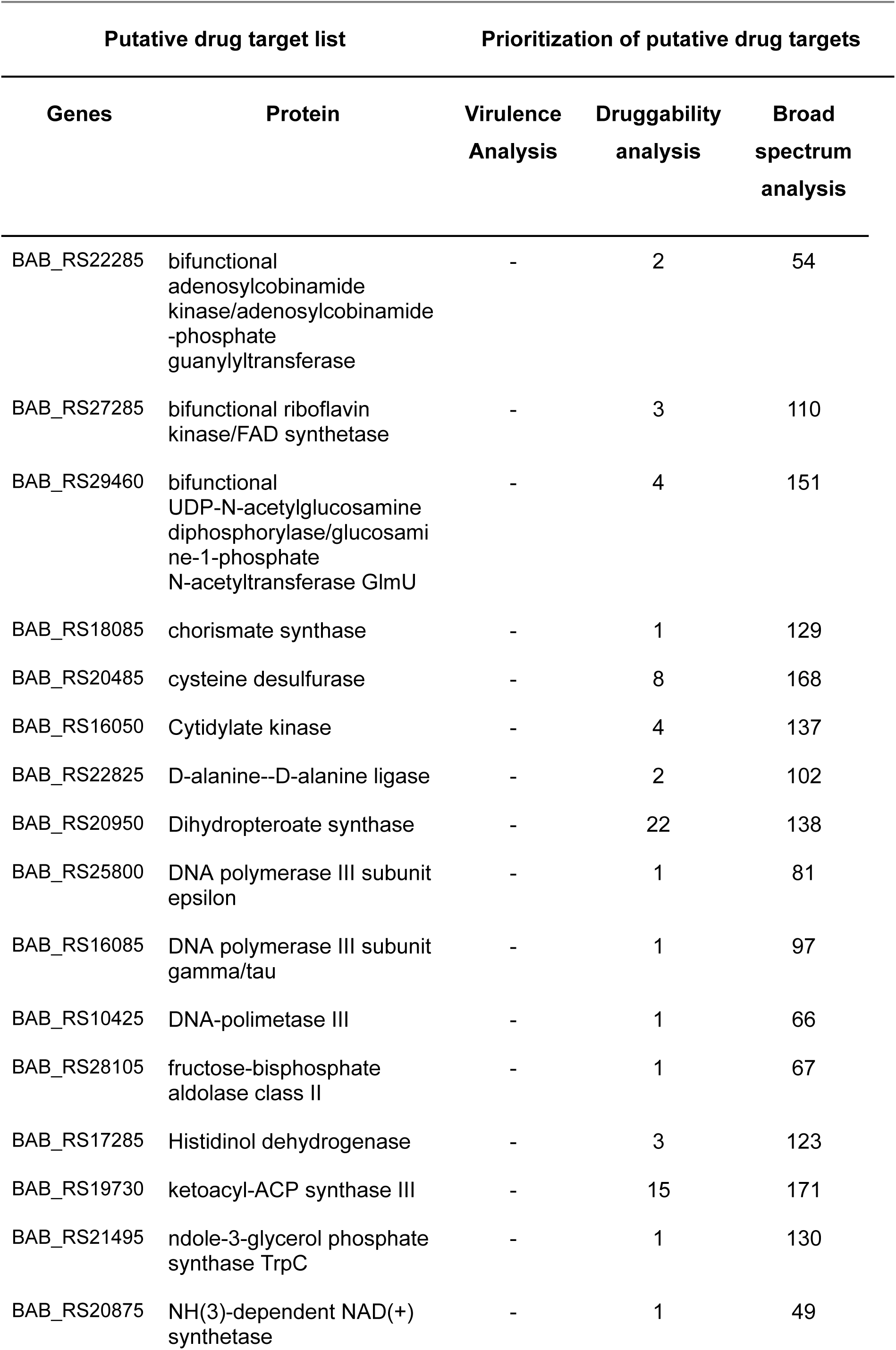
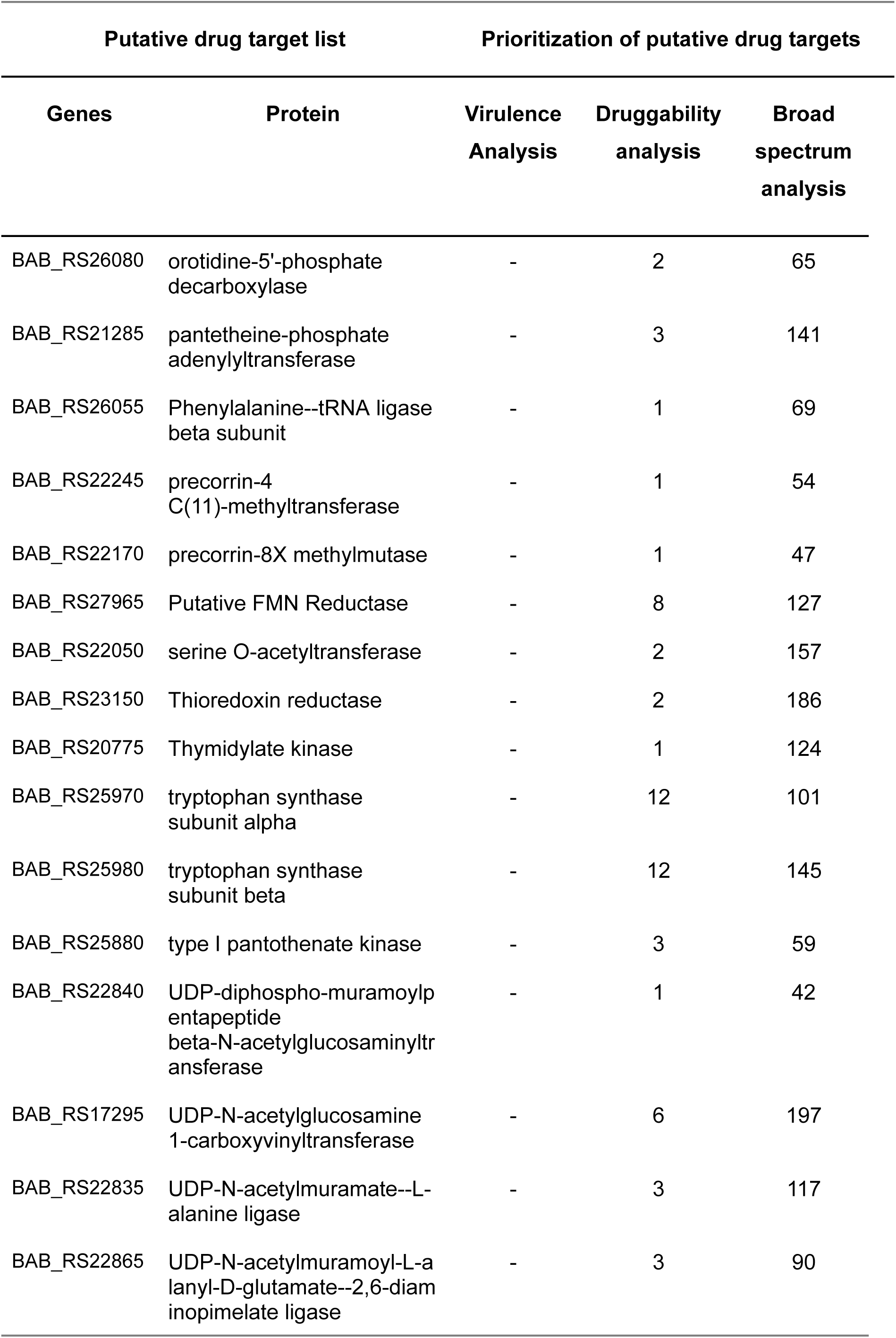
Targets predicted in at least two of three criteria tested. Only five genes passes all criteria. All essential genes and proteins along with the number of interacting drugs are listed in the table.

The virulence of *Brucella* relies on its ability to proliferate within phagocytic host cells while evading the host’s killing mechanisms. In *B. suis,* the LPS-O chain has been implicated in inhibiting early fusion between lysosomes and *B. suis* phagosomes within macrophages, thereby facilitating its intracellular survival (Cerdá et al., 1998; Porte et al., 2003). Through drug target prediction analysis, the virulence factors identified, namely *lpxA*, *lpxC*, *kdsB*, and *kdsA*, are involved in different stages of lipid A biosynthesis, a crucial component of the LPS structure. The *kdsA* gene has previously been recognized as a potential drug target in studies focused on *P. aeruginosa*, *Klebsiella pneumoniae*, and *Serratia marcescens*, all of which are opportunistic pathogens (Perumal et al., 2010; Gupta et al., 2016; Cesur et al., 2020). The *lpxC* gene has also been explored as a drug target for *P. aeruginosa* and *K. pneumoniae* (Mdluli et al., 2006; Ahmad et al., 2016), while inhibition of the *lpxA* gene product has been investigated through the development of inhibitor peptides (Jenkins and Dotson, 2012; Dangkulwanich et al., 2019). The enzyme 3-hydroxyacyl-ACP dehydratase *FabZ* was also identified as a virulence factor. Its structure, inhibitors, and inhibition mechanism have been characterized to validate its potential as a target for antimicrobial development against pathogenic bacteria (Liu et al., 2005; Zhang et al., 2008; Kumar et al., 2018). In the reconstruction of the iBP932 model, the biosynthesis pathways of lipid A and saturated fatty acids were identified as potential virulence-associated pathways, suggesting the genes *lpxA*, *lpxC*, *kdsB*, *kdsA*, and *FabZ* as potential drug targets. Currently, there is limited data exploring these targets specifically for *B. abortus.* However, a Tn-seq experiment demonstrated that five of these genes were experimentally validated as essential genes (Sternon et al., 2020).

The prediction of essential genes and analysis of non-human homologs for both HBF and MFG conditions have revealed important genes involved in vitamin biosynthesis. These genes present potential as drug targets and warrant further investigation into their impact on *B. abortus* metabolism and virulence. The final step in prioritizing drug targets involved a broad-spectrum analysis using the PIBT web browser for the identified 69 druggable proteins. Among them, a total of 59 proteins were identified as having broad-spectrum potential, with 35 of them being associated with 100 or more pathogenic bacteria. Notably, the targets *lpxC*, *kdsB*, *kdsA*, and *FabZ* met all the criteria tested and are considered the most promising broad-spectrum drug target candidates.

## Conclusion

This study developed the first *B. abortus* GEM based on genomic data and information from databases and literature. The model enabled prediction of essential genes and the simulation under different carbon and nitrogen sources. Through the application of this model, a series of analyses was performed, leading to the identification of four essential genes that could be explored as potential targets for the treatment of brucellosis and other infectious diseases, given their identification as broad-spectrum targets. This model can serve as a tool for construction of other *Brucella* species GEMs, as well as integrated metabolic models that can simulate the systemic pathogen-host interaction metabolism. The integration of other omics data, such as transcriptomics and proteomics, can contribute to the construction of context-specific models in order to simulate biological conditions more accurately. Additionally, the development of models with enzymatic constraints based on iBP932 can be utilized to conduct more precise analyses, helping to elucidate the processes of *B. abortus* intracellular pathogenesis.

## Acknowledgements

This work was supported by the Conselho Nacional de Desenvolvimento Científico e Tecnológico (CNPq), Coordenação de Aperfeiçoamento de Pessoal de Nível Superior (CAPES) and Fundação de Amparo à Pesquisa do Estado de Minas Gerais (FAPEMIG). The authors thank the Diretoria de Tecnologia de Informação (DTI) at Universidade Federal de Viçosa for availability of the computational cluster and software used in this work.

