## Supplementary material for "First Genome-Scale Metabolic Modeling of *Brucella abortus* Predicts Pathogen Gene Essentiality and New Drug Target": Supplementary File S1.docx

**Growth Rate Calculation**

The growth rate was calculated based on the growing curve present in the Viadas et al., 2010 work. The *B. abortus* cells was cultivated in a Brucella synthetic liquid medium of Gerhardt (Gerhardt., 1958). The OD values considered for the calculation was 0.7 and 0.2 and the referent time of 5 and 4 days converted to hours. The calculation was made considering the linear phase, referent to the log phase of the bacterial growth using the following formula:

μ = ln 0.7 -ln 0.2	= 0.052 h^-1^

_____________

120 - 96

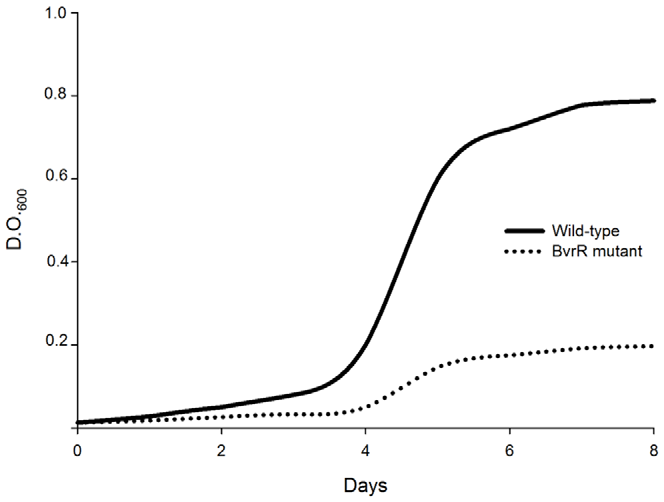

Figure 3 referent to Viadas et al., 2010 work of *B. abortus* growth in synthetic minimal media.
